# Re-Identification of Individuals in Genomic Data-Sharing Beacons via Allele Inference

**DOI:** 10.1101/200147

**Authors:** Nora von Thenen, Erman Ayday, A. Ercument Cicek

**Affiliations:** Computer Engineering Department, Bilkent University, 06800 Bilkent, Ankara, Turkey; Computational Biology Department, Carnegie Mellon University, 5000 Forbes Ave., Pittsburgh, PA 15213

## Abstract

Genomic datasets are often associated with sensitive phenotypes. Therefore, the leak of membership information is a major privacy risk. Genomic beacons aim to provide a secure, easy to implement, and standardized interface for data sharing by only allowing yes/no queries on the presence of specific alleles in the dataset. Previously deemed secure against re-identification attacks, beacons were shown to be vulnerable despite their stringent policy. Recent studies have demonstrated that it is possible to determine whether the victim is in the dataset, by repeatedly querying the beacon for his/her single nucleotide polymorphisms (SNPs). In this work, we propose a novel re-identification attack and show that the privacy risk is more serious than previously thought. Using the proposed attack, even if the victim systematically hides informative SNPs (i.e., SNPs with very low minor allele frequency -MAF-), it is possible to infer the alleles at positions of interest as well as the beacon query results with very high confidence. Our method is based on the fact that alleles at different loci are not necessarily independent. We use the linkage disequilibrium and a high-order Markov chain-based algorithm for the inference. We show that in a simulated beacon with 65 individuals from the CEU population, we can infer membership of individuals with 95% confidence with only 5 queries, even when SNPs with MAF less than 0.05 are hidden. This means, we need less than 0.5% of the number of queries that existing works require, to determine beacon membership under the same conditions. We further show that countermeasures such as hiding certain parts of the genome or setting a query budget for the user would fail to protect the privacy of the participants under our adversary model.

## 1 Introduction

Exciting times are on the horizon for the genomics field with the announcement of the precision medicine initiative [4] which was followed by the $55 million funding by NIH for the sequencing of a million individuals and AstraZeneca’s project of sequencing two million individuals [9]. Even though such million-sized genomic datasets are invaluable resources for research, sharing the data is a big challenge due to re-identification risk. Several studies in the last decade have shown that removal of personal identifiers from genomic data is not enough and that individuals can be re-identified using allele frequency information [6, 12, 8, 15, 3].

Genomic data-sharing beacons (referred to as beacons from now on) are the gateways that let users and data owners exchange information without -in theory-disclosing any personal information. A user who wants to apply for access to the dataset can learn whether individuals with specific alleles of interest are present in the beacon through an online interface. More specifically, the user submits a query, asking whether a genome exists in the beacon with a certain nucleotide at a certain position, and the beacon answers “yes” or “no“. Beacons are easy to set up systems that provide very restricted access to the stored data. The Beacon Project is an initiative by the Global Alliance for Genomics and Health (GA4GH) which creates policies to ensure standardized and secure sharing of genomic data.

Beacons were considered safe as allele frequencies are not involved in the query result and the binary answers for allele presence seem far from being informative for an attack. However, in 2015, Shringarpure and Bustamante introduced a likelihood-ratio test (LRT) that predicts if an individual is in the beacon or not, by repeatedly querying the beacon for SNPs of the victim (dubbed the SB attack) [13]. The method does not use the allele frequencies and can compensate sequencing errors. They show that they could re-identify an individual in a beacon with 65 European individuals from the 1000 Genomes Project [14] with 250 queries (with 95% confidence). In their scheme, both the queries posed and the answers received from the beacon are assumed to be independent, therefore the hypothesis is tested based on a binomial test. Very recently, the work by Raisaro *et al.* showed that if the attacker has access to the MAFs of the population, s/he can sort the victim’s SNPs and query the SNPs starting from the one with the lowest MAF (dubbed the Optimal attack) [10]. Unlike the SB attack, queries are not random in this case. As low MAF SNPs are more informative, Raisaro *et al.* show that fewer queries are needed to re-identify an individual. Furthermore, Raisaro *et al.* proposed countermeasures against re-identification attacks such as adding noise to the beacon results and assigning a budget to beacon members which limits the number of informative queries that can be asked on each member.

In this paper, we introduce two new inference-based attacks that (i) carefully select the SNPs to be queried and predict query results of the beacon, and (ii) infer hidden or missing alleles of a victim’s genome. First, we show that if the queried locus is in linkage disequilibrium ^1^ with others, it is enough to query for that particular allele, as the attacker can infer the answers of the other alleles with high confidence [7]. We refer to this method as the QI-attack (query inference attack). Second, we introduce the GI-attack (genome inference attack) which recovers hidden parts of a victim’s genome by using a high-order Markov chain [11].

We show that in a simulated beacon with 65 European individuals (CEU) from the HapMap Project [5], our QI-attack requires 282 queries and our GI-attack requires only 5 queries on average to re-identify an individual, whereas the SB attack requires 19,525 queries and the Optimal attack requires 415 queries, all at the 95% confidence level when the victim’s SNPs with MAFs *<* 0.03 are hidden. Therefore, the attacker models presented here can efficiently work when certain regions in the genome of the victim are systematically hidden as a security countermeasure. The number of queries required by the SB and the Optimal attacks substantially increase as more SNPs are concealed, while the GI-attack still requires only a few queries on average. Finally, we show that the QI-attack can still re-identify individuals despite the stringent query budget countermeasure proposed by [10] and the beacon censorship countermeasure proposed by [13].

We demonstrate that the beacons are more vulnerable than previously thought and that the proposed countermeasures in the literature still fail to protect the privacy of the individuals. The contributions of this paper can be summarized as follows:

– By inferring query results and alleles at certain positions, we show that it is possible to significantly decrease the number of required queries compared to other attacks in the literature [13, 10].
– We show that beacons are vulnerable even under a weaker adversary model, in which informative parts of a victim’s genome are concealed (such as all SNPs with an MAF less than a threshold).
– We discuss the feasibility and the effectiveness of the proposed countermeasures in the literature and show that using the presented attack models, the participants are still under risk.

The rest of the manuscript is organized as follows: We describe the methods in Section 2 and then present the results in Section 3. Section 4 discusses the results and the effectiveness of countermeasures proposed in the literature. Finally, we conclude in Section 5.

## 2 Material and Methods

In this section, we first describe attacker models in the literature (i.e. SB attack [13] and the Optimal attack [10]), and then describe our proposed attacks. In our first proposed model, the attacker not only has access to MAFs of the victim’s population, but also can access or calculate the corresponding linkage disequilibrium values from public resources (QI-attack). In the second model, the attacker has the same background knowledge as the QI-attack, and also has access to VCF files of people from the victim’s population from public sources (GI-attack). The four different attacker models (SB attack [13], Optimal attack [10], QI-attack, and GI-attack) are described in Figure 1. We consider two scenarios. Scenario 1 assumes the attacker has access to the full genome of the victim ^2^.Scenario 2 considers a more realistic and weaker attacker model. As publicly available genomic data is typically found partially, in this scenario, some SNPs are systematically hidden. That is, SNPs with MAF *< t* are not available to the attacker.

### 2.1 Background: SB attack & Optimal attack

Shringarpure and Bustamante proposed the SB attack, which queries a beacon for the victim’s heterozygous SNP positions. Queried SNPs are picked randomly and a likelihood ratio test (LRT) statistic is calculated. The null hypothesis (*H*_0_) refers to the query genome not being in the beacon. Under the alternative hypothesis (*H*_1_), the query genome is a member of the beacon. The attacker model is visualized in Figure 2(a). The log-likelihood under the null hypothesis has been defined as

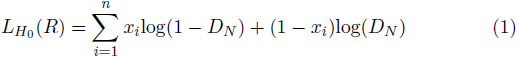

where *R* is the response set and *D*_*N*_ the probability that no individual in the beacon has the queried allele at that position. *x*_*i*_ is the answer of the beacon to the query at position *i* (1 for yes, 0 for no), and *n* is the total number of posed queries. Accordingly, the log-likelihood of the alternative hypothesis has been stated as

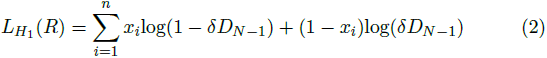

where *D*_*N-*1_ represents the probability of no individual except for the queried person having the queried SNP. represents a possible sequencing error. Finally, the LRT statistic is stated as follows:

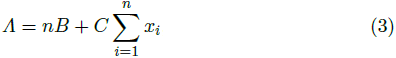

where *B* and *C* are defined as *B* = log(*D*_*N*_ */δD*_*N-*1_) and *C* = log(*δD*_*N-*1_(1 *- D*_*N*_ */D*_*N*_ (1 *-δD*_*N-*1_)), respectively. The null hypothesis is rejected for any *Λ* that is less than a certain threshold.

**Fig. 1.**
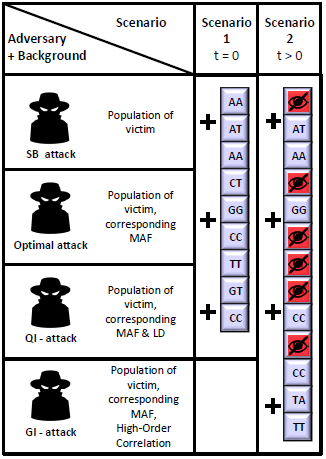
Four attacker models: SB attack [13], Optimal attack [10], QI-attack, and GI-attack and their background knowledge for two scenarios are shown. In the first scenario *t* = 0 and in the second scenario *t >*0, where *t* is the threshold up to which SNPs of the victim with an MAF *< t* are hidden as a countermeasure. In Scenario 1, the attacker has access to the full genome of the victim (no hidden SNPs). In Scenario 2, SNPs with an MAF *< t* are hidden and the attacker has partial access to the genome of the victim.

The Optimal attack introduced by Raisaro *et al.* integrates publicly available MAF information into the attacker’s background knowledge [10]. In this attack, the victim’s SNPs are sorted with respect to their MAFs. The beacon is queried starting from the first heterozygous SNP with the lowest MAF. The model of this attack is illustrated in Figure 2(b). In this setting, the computations of *D*_*N-*1_ and *D*_*N*_ depend on the queried position *i* and change at each query as shown as follows:

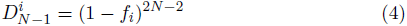

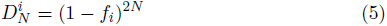

where *f*_*i*_ represents the MAF of the SNP at position *i*. Accordingly,Λ changes as follows:

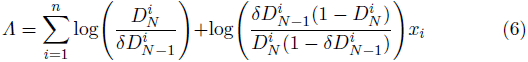

### 2.2 Query Inference Attack

The QI-attack uses pairwise SNP correlations (LD) in order to infer the answers of unasked queries from previously answered queries. In this model, the attacker uses the LD value of a SNP pair to calculate the correlation of two minor alleles at the corresponding loci. The correlation is equal to the probability of the two minor alleles occurring together. Let *p*_2_ be the MAF of SNP *A* (with minor allele *a*) and *q*_2_ be the MAF of SNP *B* (with minor allele *b*). Assuming *A* and *B* are in LD, the probability of two major or two minor alleles in these loci occurring together increases. This can be calculated as *P r*(*ab*) = *p*_2_*q*_2_ + *D*, where *D* represents the strength of the correlation of the two SNPs (see Appendix A for details). On this basis, the attacker constructs a SNP network that uses weighted, directed edges between SNPs in high LD (see Figure A.1 in Appendix A). The weight corresponds to the probability of two minor alleles occurring together. Figure 2(c) illustrates this model. First, the attacker selects the SNPs to be queried. This step is identical to the Optimal attack and leads to a set of candidate SNPs *S* to be queried, starting from the lowest MAF *SNP*_*i*_. Second, if any non-queried *SNP*_*j*_ in *S* is a neighbor of *SNP*_*i*_ in the SNP network, the attacker infers the result of the query and does not pose a query for *SNP*_*j*_. In the following, we present the null and the alternative hypotheses in this model which also integrates the inference error.

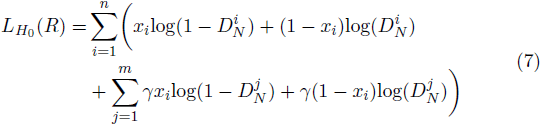

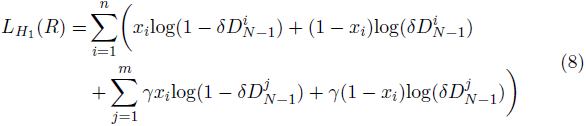

where *n* is the number of posed queries, *m* is the number of neighbors that can be inferred for each posed query *x*_*i*_, and γ corresponds to the confidence of the inferred answer, obtained from the SNP network. *Λ* is then determined as

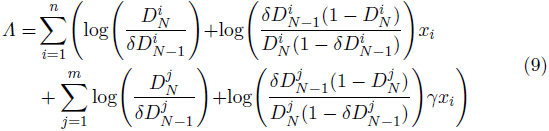

By not querying the beacon for answers that can be inferred with high confidence, this model requires less number of queries compared to the Optimal attack, while achieving the same response set. For more detail, see Appendix B.

**Fig. 2.**
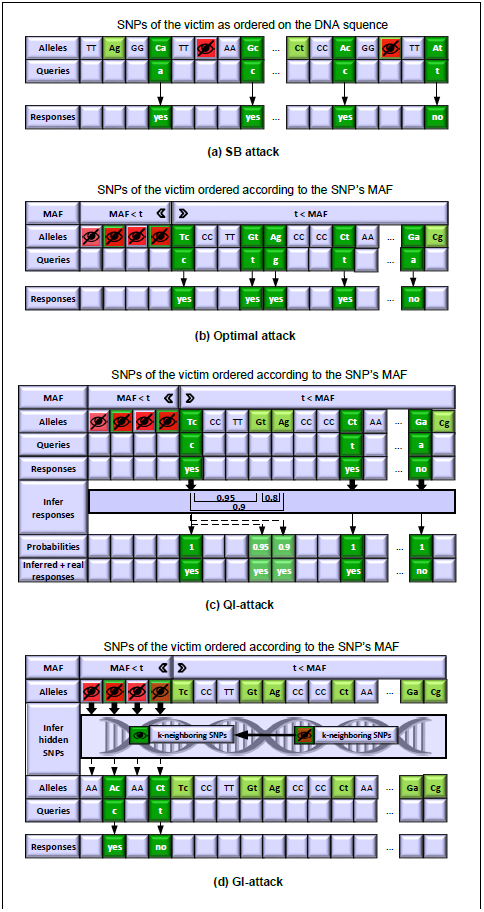
System models of the four attacker models (a) SB attack [13], (b) Optimal attack [10], (c) QI-attack and (d) GI-attack. Upper-case letters represent the major allele at a SNP position and the lower-case letters the corresponding minor allele. The SB attack randomly selects the minor allele from heterozygous SNP positions of the victim and queries those. The Optimal attack first sorts the heterozygous SNPs regarding their MAFs and queries for the minor alleles starting with the lowest frequency. Depending on the threshold *t*, SNPs with MAF *< t* are hidden and are not available to the attacker. The QI-attack extends the Optimal attack by inferring beacon answers using LD correlations between SNP pairs. The GI-attack infers the hidden SNPs with MAFs *< t*, using a high-order Markov chain and queries the beacon for the minor alleles of those positions.

### 2.3 Genome Inference Attack

Individuals may publicly share their genomes by taking necessary precautions, such as hiding their sensitive SNP positions with MAFs *< t* (Scenario 2 in Figure 1). The GI-attack performs allele inference to recover hidden SNP positions and infers alleles at the victim’s hidden loci. Note that Scenario 1 (in Figure 1) is not applicable to the GI-attack, since in that scenario, the attacker can access SNPs with low MAFs. The attacker uses a high-order Markov chain to model SNP correlations as described by Samani *et al.* [11].

The model of this attack is illustrated in Figure 2(d). Depending on the threshold *t*, the attacker infers SNP positions with MAF *< t* that are not available in the victim’s VCF file. Based on the victim’s genome sequence, the attacker calculates the likelihood of the victim having a heterozygous position at the chosen SNP position *i* as follows

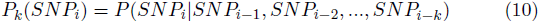

where *k* is the order of the Markov chain. In order to use a high-order Markov chain to infer hidden SNPs, genome sequences from public sources such as the 1000 Genomes project or HapMap can be used to train the model. ^3^ Accordingly, Samani *et al.* define the *k* ^*th*^-order model as

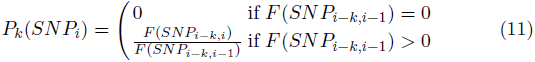

where *F* (*SNP*_*i,j*_) is the frequency of occurrence of the sequence that contains *SNP*_*i*_ to *SNP*_*j*_. The SNPs are ordered according to their physical position on the genome. The model works by comparing the SNPs in *SNP*_*i,j*_ which are prior to *SNP*_*i*_ on the genome sequence to the same SNP positions in the training dataset. If the training set contains other genomes with the same SNP sequence and these sequences are followed by a heterozygous position, we can calculate the probability of *SNP*_*i*_ being heterozygous for our victim. As an example, the victim’s 4 ^*th*^-order SNP sequence is [AA, AT, CC, TT]. We would now like to determine whether the following *SNP*_*i*_, that is hidden in the VCF file at hand, is likely to be a heterozygous position. Therefore, we identify other genomes in the training dataset with the same sequence and compute the frequency of this sequence being followed by a heterozygous position. That is, [AA, AT, CC, TT] *→* [AG]. As a result, we can determine the probability of the four SNPs being followed by a heterozygous position, which we can use to query the beacon.

If the calculated likelihood of the victim having a heterozygous position is high enough (in this case equal to 1), the attacker queries the beacon for the inferred SNP position, starting from the SNP with the lowest MAF.

## 3 Results

To evaluate our attacks, we tested our methods on (i) a simulated beacon and compared our results with the SB attack [13] and the Optimal attack [10] (Section 3.1), and (ii) the beacons of the beacon-network ^4^ operated by GA4GH Beacon Network and compared our results with the Optimal attack [10] (Section 3.2).

### 3.1 Re-identification on a Simulated Beacon

**Fig. 3.**
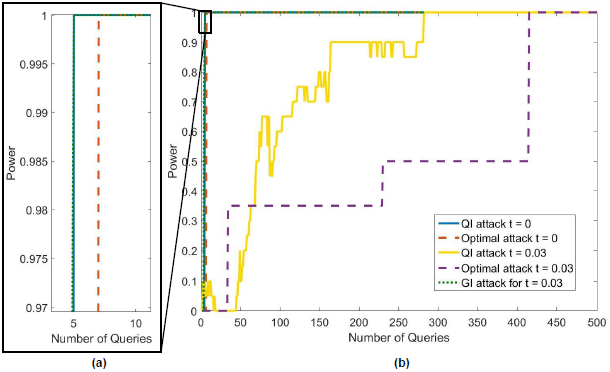
(a) Close-up of the power curves, where number of queries *<* 10. (b) Power curves of the Optimal attack [10], the QI-attack, and the GI-attack for different thresholds of *t* on a beacon with 65 members constructed with individuals from the CEU dataset of the HapMap project. *t* indicates the threshold up to which SNPs with an MAF *< t* are hidden as a countermeasure.

In this section, we evaluated the performance of the four attacks on a simulated beacon with 65 people from the CEU population of the HapMap dataset. While testing for the alternative hypothesis, we used 20 randomly-picked people from the beacon. For the null hypothesis, we used 40 additional people from the same population of the HapMap project. The CEU population is the population of choice because previous works (SB attack [13] and Optimal attack [10]) have also been evaluated on this population. The LD scores, allele frequencies, and genotype data were also obtained from the CEU dataset of the HapMap project [5]. For the GI-attack, we used a 4 ^*th*^-order Markov chain (see Appendix C for details of selecting the order).

**Table 1.** Average number of queries needed to receive the first negative response for the SB attack [13], the Optimal attack [10], the QI-attack, and the GI-attack for different thresholds of *t* on a beacon with 65 members constructed with 40 case individuals from the CEU dataset of the HapMap project. *t* indicates the threshold up to which SNPs with an MAF *< t* are hidden. As the GI-attack concentrates on inferring hidden parts of the genome, we do not consider *t* = 0 (nothing is hidden) for the GI-attack.

|  | # of queries |  |  |  |
| --- | --- | --- | --- | --- |
| $t$ | SB attack | Optimal attack | QI-attack | GI-attack |
| 0 | 1,418 | 3 | 3 | NA |
| 0.03 | 19,525 | 270 | 160 | 2 |
| 0.05 | 56,759 | 1,495 | 1,031 | 2 |

We show the power curves for the Optimal, the QI-attack and the GI-attack each at 5% false positive rate in Figure 3 and the number of queries needed to receive the first negative response in Table 1. We empirically build the null hypothesis. That is, we determine the distribution of *Λ* under the null hypothesis using the 40 people who are not in the beacon. When *Λ* is less than a threshold, the null hypothesis is rejected. Similar to Raisaro *et al.* [10], we reject the null hypothesis when *Λ< tα*. We find the threshold *tα* from the null hypothesis with *α* = 0.05 (corresponding to 5% false positive rate). The power 1 − *β* is then the proportion of the individuals in the control set having a *Λ* value, where *Λ< tα*. See Appendix D for more information on the power calculation.

We observed that the SB attack requires the highest number of queries (1,400 - 56,800). The QI-attack requires 30% less number of queries on average compared to the Optimal attack. The GI-attack requires only 5 queries for all tested thresholds of *t*.

Compared to the monotonically increasing behavior of the power curves for the Optimal attack, the power curve for the QI-attack shows a zig-zag behavior. This is because *t*_*α*_is recalculated at each posed query and the *Λ* values change based on the number of inferred queries.

The threshold *t* of hidden SNPs significantly affects the performance of the attacks. As *t* increases, more common SNPs are available to the attacker which means that the likelihood of another individual in the beacon having the same allele increases. When the beacon was queried for each of the 40 people who are not in the beacon, the SB attack was not able to receive a “no” response with 100,000 queries, (i) for 4 people when SNPs with an MAF *<* 0.04 were hidden and (ii) for 12 people when SNPs with an MAF *<* 0.05 were hidden. Therefore, it was not possible to correctly determine beacon membership for all test individuals to reach 100% power for larger *t* values. Compared to the GI-attack, the Optimal and the QI-attack required a significantly higher amount of queries to determine beacon membership and reach 100% power. The GI-attack successfully determined the correct status for all 40 individuals despite the high threshold of *t* with only a few queries.

### 3.2 Re-identification on Existing Beacons

We tested our methods on the beacons of the beacon-network. We selected an individual from the Personal Genomes Project (PGP) (Person’s id: PGP180/hu2D53F2) [2] as the victim. To determine if this person is a member of the beacons, we applied the SB attack as ground truth as detailed in Appendix E. For the QI-attack, we used the same SNP network as for the simulated beacon in Section 3.1 (based on the CEU population of HapMap). The Markov chain of the GI-attack was trained on the CEU population of the HapMap [5] dataset. We again used a 4 ^*th*^-order Markov chain (see Appendix C for details of selecting the order).

The beacons can return an empty response, that is, the beacon has no information at that position, a “no“-response, and a “yes“-response.We consider two cases for the evaluation of the query results. In the first case, an empty answer is treated as a “no” (results shown in Table 3.2), in the second case an empty answer is not treated as a “no“, as it is also possible that the beacon has a different copy of the victim’s genome (results shown in Table F.1 in Appendix F). As the results are similar, we concentrate on the first case in the following.

Unlike all other beacons, the 1000 Genome Project beacon required fewer number of queries for re-identification as *t* is increased. Note that the victim’s SNPs are sorted based on the CEU population’s allele frequencies. Thus, SNPs that we query are not necessarily the rarest in the queried beacon, which can explain this behavior. Furthermore, the SNP network used is also based on the CEU population and therefore, does not include all SNPs of the victim’s genome.

The GI-attack performed as expected, that is constant over the two tested thresholds of *t* and outperformed the Optimal attack [10] as well as the QI-attack for *t >* 0. For the 1000 Genomes Beacon the GI-attack required the same amount of queries as the other attacks, as the number of queries needed are already very low.

**Table 2.** Number of queries required to receive a “no” within 1000 queries to existing beacons using an individual from PGP [2] when *t* = {0, 0.03, 0.05} for the Optimal attack [10], the QI-attack, and the GI-attack. Here, empty answers (i.e., the beacon has no information about the queried locus in the underlying dataset and returns neither a “no” nor a “yes“) are not considered as a “no” response. “-” means in no “no” was found in 1000 queries.

| Beacon Name | $t =$ | | |
| --- | --- | --- | --- |
|  | Optimal attack<br>0\|0.03\|0.05 | QI-attack<br>0\|0.03\|0.05 | GI-attack<br>0.03\|0.05 |
| Known VARIants | -\|-\|- | -\|-\|- | -\|- |
| Broad Institute | 2\|2\|2 | 2\|2\|2 | 1\|1 |
| 1000 Genomes Project | 4\|3\|2 | 4\|3\|2 | 3\|3 |
| Cafe CardioKit | -\|-\|- | -\|-\|- | -\|- |
| Wellcome Trust<br>Sanger Institute | 1\|1\|1 | 1\|1\|1 | 1\|1 |
| NCBI | -\|-\|- | -\|-\|- | -\|- |
| ICGC | 1\|-\|- | 1\|-\|- | 1\|1 |
| AMPLab | 20\|45\|73 | 20\|40\|73 | 39\|39 |
| 1000 Genomes<br>Project phase 3 | 20\|130\|250 | 20\|116\|250 | 48\|48 |

In summary, for 6 of the 9 tested beacons, we were able to determine that the victim is not a member of the beacons. For the Known VARiants (Kaviar), the Cafe CardioKit, and the NCBI, it was not possible within 1,000 queries (Figure 3.2). Overall, we observed that the experiments on real beacon support our findings in Section 3.1. That is, the Optimal and the QI-attack need more queries as *t* increases, the GI-attack is stable over all thresholds, and the QI-attack requires less queries than the Optimal attack.

## 4 Discussion

Recent works by Shringarpure and Bustamante [13] and Raisero *et al.* [10] have shown, that beacon servers fail at protecting their members’ privacy. As beacons are often associated with a certain phenotype, the membership identification of an individual could leak sensitive information. They proposed countermeasures such as (i) user budget, (ii) adding noise, and (iii) increasing beacon size to improve the security level of existing beacons.

In this work, we have shown that beacon membership can be detected with even a lower number of queries and with high confidence, despite strict countermeasures. Overcoming the proposed countermeasures is possible by including publicly available information such as MAF, LD, and VCF files (from e.g., HapMap [5] or 1000 Genomes Project [14]) into the attacker model. Previous works in the field of genomics and privacy have shown that it is possible to increase the success rate of genomic re-identification attacks by including LD information into the attacker model. Namely, Wang *et al.* showed in 2009 that individuals can be reidentified by using (i) publicly available SNP-to-disease correlation information, and (ii) SNPs in LD [16]. In 2013, Humbert *et al.* showed how LD can be used to build a framework to reconstruct the genomes of people using the genome of a family member [7].

The success of our QI-attack depends significantly on the structure of the underlying SNP network. The larger and denser the network becomes, the more query responses can be inferred. Additionally, the strength of the SNP correlations is an important factor. In this work, we included SNP pairs that are in strong LD (i.e. *r* ^2^ *>* 0.7) in our SNP network to limit inference error.

The GI-attack shows that even if genomes do not contain any SNPs with low MAFs, individuals’ privacy is not ensured, as it is possible to infer these loci using information from publicly available datasets (e.g., HapMap [5] or 1000 Genomes Project [14]). As shown in Appendix G, the GI-attack still performs as good even when the attacker trains the high-order Markov chain on a different population than the victim’s.

Our experiments on a simulated beacon (Section 3.1) and existing beacons (Section 3.2) show that as the threshold up to which SNPs of the victim with an MAF *< t* are hidden (*t*) increases, our attacks require fewer queries than existing attacks (SB attack [13] and Optimal attack [10]). Table 3.2 shows that for the existing beacons the number of queries needed increases as *t* increases and that the margins are even larger compared to the simulated beacon (Table 1).

Shringarpure and Bustamante discussed different countermeasures, such as (i) increasing the beacon size, (ii) sharing only small genomic regions, (iii) using single population beacons, (iv) not publishing the metadata of a beacon, and (v) adding control samples to the beacon dataset [13]. Lately, Aziz *et al.* proposed two algorithms which are based on randomizing the response set of the beacons with the goal of protecting beacon members’ privacy while maintaining the efficacy of the beacon servers [1]. Raisaro *et al.* have analyzed the behavior of the beacon when applying three different countermeasures [10]. First, they propose the beacon should only respond “yes” for an allele if multiple samples have it. The second countermeasure adds noise to the responses. However, this countermeasure significantly reduces the utility of the dataset and is unacceptable for researchers working on data-sharing beacons. Instead, the beacon could return an empty answer. Final countermeasure is assigning a query budget per sample. That is, every member of the beacon is assigned with a certain budget that is reduced if a query to the beacon matches the sample. As an example, if a user queries the beacon for allele A in position 1000 of chromosome 21, then the budget of every member with an allele A in that position is reduced. The amount of the budget reduction is determined based on the risk of the query, where the lower the allele frequency of the queried allele is, the higher the risk becomes. The budget is calculated as *b*_*i*_ = log(*p*), where Raisaro *et al.* use *p* = 0.05. The risk then is calculated as 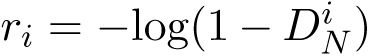 If the budget of a beacon member is depleted, the beacon stops including the member into the beacon responses.

An attacker using the QI-attack can overcome this countermeasure. For instance, in our simulated beacon as described in Section 3.1, an attacker using the Optimal attack needs 7 queries to re-identify the victim (individual “NA12272” of the HapMap project [5]), when no SNPs are hidden. However, the beacon would start giving false responses after 6 queries as the budget would be depleted, which means the attack would fail. By using the QI-attack, an attacker would only need 5 queries. Therefore, a query budget that is merely based on the SNPs’ MAFs and that does not consider SNP correlations would fail to protect an individual’s privacy. An attacker using the QI-attack would not exhaust the budget but still be able to determine the victim’s beacon membership.

As the countermeasures of the literature fail to protect beacon members privacy, we will concentrate on developing new countermeasures based on SNP correlations and allele frequencies as future work.

## 5 Conclusion

Throughout the course of this work, we showed that data-sharing beacons are sensitive to re-identification attacks. Additionally, we showed that countermeasures that do not consider the MAFs and correlations of SNPs fail to protect the beacon members’ privacy. Furthermore, even if individuals apply countermeasures before releasing their genome, such as systematically hiding SNPs with low MAFs, their privacy still could be at stake. Therefore, new countermeasures are needed to ensure privacy of individuals.

## Appendices

### A SNP Network

When two SNPs *A* and *B* are in LD, the probability of two major or two minor alleles occurring together increases or decreases by *D*. As shown in Table A.1 this leads to the formula: *P r*(*ab*) = *p*_2_*q*_2_ + *D*, where *p*_2_ is the minor allele frequency of SNP *A* (with minor allele *a*) and *q*_2_ is the minor allele frequency of SNP *B* (with minor allele *b*). *D* is calculated as follows: 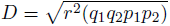 where *q*_1_ and *p*_1_ are the major allele frequencies and *r* ^2^ is a common measure of LD. To determine whether the LD correlation increases or decreases the probability of the two loci occurring together *D* ^*'*^ is needed. *D* ^*'*^ 0.5 implies *D* is added, whereas *D* ^*'*^ *<* 0.5 leads to a subtraction of *D*.

**Table A.1.** Relationship between Linkage Disequilibrium (LD) measured by *D* between the SNPs *A* and *B* and their allele frequencies.

| | $Pr(A) = p_1$ | $Pr(a) = p_2$ |
| --- | --- | --- |
| $Pr(B) = q_1$ | $p_1q_1 + D$ | $p_2q_1 - D$ |
| $Pr(b) = q_2$ | $p_1q_2 - D$ | $p_2q_2 + D$ |

The SNP network that is used to determine correlated SNPs is build as a directed graph. The edges are labeled with the probability of two minor alleles occurring together in the considered SNP positions as shown in the lower right field of Table A.1. In order to ensure high correlation between the SNPs, only LD relationships between SNP pairs with an *r* ^2^ value of more than 0.7 were considered. The average of the correlation between SNP pairs and therefore the labels on the edges of the SNP network is 0.9511. An example of a SNP network with 5 nodes is shown in Figure A.1.

**Fig. A.1.**
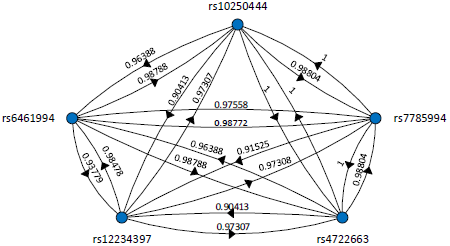
A SNP network that contains 5 nodes (i.e., SNPs). The SNP network is adirected graph, where the weight of edges correspond to the correlation between SNPs.No edge between a pair of SNPs means the correlation is less than the r2 threshold.The correlation is in the most cases not symmetric, since it depends on the minor allelefrequencies of the SNP pair. This example shows a fully connected graph, which is not necessarily the case for all SNP networks.

### B LRT - Query Inference Attack

The likelihood of the null hypothesis is calculated as follows:

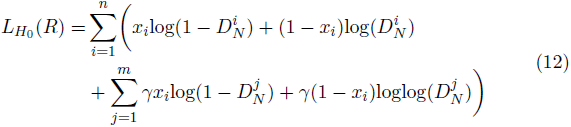

where, *x*_*i*_ is the response of queried *SNP*_*i*_, *n* the number of posed queries, *m* the number of queries that were inferred with query *SNP*_*i*_, and γ the confidence of the inferred responses.

Accordingly, the likelihood of the alternative hypothesis is found as follows:

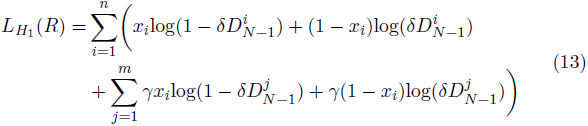

Using (12) and (13), *Λ* can be calculated as shown in Equation 14.

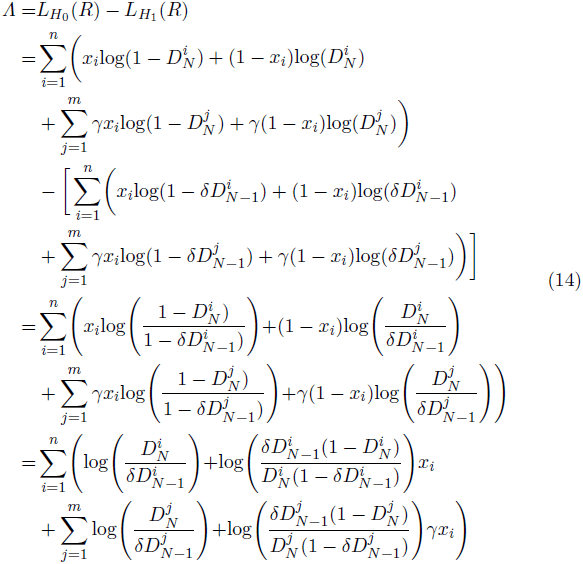

### C High-Order Markov Chain

Individuals may hide certain loci on their genome before publishing their VCF files. It is possible to infer these hidden positions by applying a high-order Markov chain as introduced by Samani *et al.*, 2015 [11]. The probability of a certain allele occurring at a specific position can then be determined as

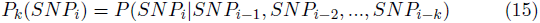

where *k* is the order of the Markov chain. Accordingly, Samani *et al.*, 2015 define the *k* ^*th*^-order model by:

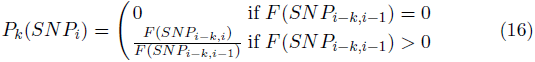

where *F* (*SNP*_*i,j*_) is the frequency of occurrence of the sequence that contains *SNP*_*i*_ to *SNP*_*j*_.

To build a high-order Markov chain to infer hidden SNPs, genome sequences from public sources such as the 1000 Genomes project or HapMap can be used to train the model. Our Markov chain is build of 100 individuals of the CEU population HapMap dataset. The SNPs are represented as 0, 1, or 2 depending on the number of minor alleles at the specific position of the genome. That is, major homozygous, heterozygous, and minor homozygous, respectively. If the VCF file of the victim has too much missing data we cannot infer beacon membership of the victim. That is, there is not enough SNP information to build the necessary *k*-order Markov chain to infer the hidden SNPs with low MAFs. Missing data can occur due to sequencing errors or can be intentionally hidden by the victim.

**Table C.1.** Comparison of different values for *k* (order of the high-order Markov chain). # of same markers shows how many markers that were inferred by the Markov chain were also asked in the Optimal attack. Distance to real response shows the amount of queries the inferred response differs from the Optimal attack’s response (on average). # of people not inferred shows the amount of people that could not be inferred for that *k*.

| $k$ (order) | # of same markers | distance to real response | # of people not inferred |
| --- | --- | --- | --- |
| 3 | 13 | 0.6 | 2 |
| 4 | 15 | 0.62 | 1 |
| 5 | 14 | 0.79 | 5 |

In order to select *k*, we considered three criteria: (i) the number of markers that are inferred by the GI-attack and also asked by the Optimal attack, (ii) the euclidean distance between the number of queries needed by the Optimal attack and the GI-attack for all tested individuals, and (iii) the number of people whose SNPs could not be inferred due to missing data.

As shown in Table C.1, we picked *k* = 4, which provides the largest number of markers, with minimum number of missed people. The distance to the real response is similar to *k* = 3 and is less than the case when *k* = 5. As our Markov chain is build on 100 individuals, we determined *k* = 5 as the maximum order to be tested to prevent over-fitting.

### D LRT - Power Calculation

The power (1 *- β*) of the LRT is determined as the proportion of control individuals (that are in the beacon) for which we can reject the null hypothesis when *Λ< t*_*α*_. The threshold *t*_*α*_ is found by building the null hypothesis with the 40 case individuals (that are not in the beacon), where *α* = 0.05 (corresponding to 5% false positive rate).

For each individual and query *x*_*i*_, we calculate the value of *Λ*, where *Λ* changes according to the attack being performed. As *t* increases, the power of the QI-attack shows a zig-zag behavior unlike the Optimal attack and the GI-attack. That is because as *t* increases, more queries are needed to determine beacon membership, and more SNPs are in inferred in the QI-attack. The more neighbors a posed query can infer from the SNP network, the more extreme the value of *Λ* changes.

Figure D.1 shows, for three example case and three example control individuals, how Ʌ steadily decreases for control individuals and clearly increases for “no” responses of case individuals (i.e., at queries 24, 26 and 84) for the Optimal attack. Here,; Ʌ decreases by a similar value for all individuals that receive a “yes” response, as only one query is asked and the queries have similar MAFs. Therefore, if Control 1 had a lower ;Ʌ value at query *x*_10_ than Case 2, Case 2 will not have a lower value than Control 1 for the following queries, unless Control 1 receives a “no” response (which leads to a significant increase in ;Ʌ but is highly unlikely for an individual in the control set).

On the contrary, Figure D.2 shows an irregular behavior of; Ʌ that is ;Ʌ does not steadily decrease unlike the Optimal attack in Figure D.1. This can be explained by the different amount of neighbors in the SNP network that can be inferred at the different loci. Considering Control 1 and Case 2 again, Control 1 can have a lower ;Ʌ value than Case 2 for query position *x*_10_. Nevertheless, if query *x*_11_ of Case 2 has a high number of neighbors to be inferred from *x*_11_ and the inferred responses are all “yes” responses, the ;Ʌ value of Case 2 decreases significantly and is now lower than the ;Ʌ value of Control 1 for *x*_11_, as *x*_11_ of Control 1 has fewer neighbors in the SNP network.

**Fig. D.1.**
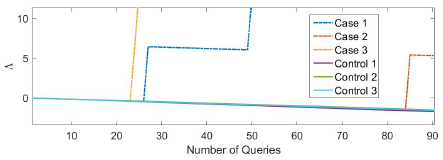
Example; Ʌ distributions for 3 of the 40 case and 3 of the 20 control individuals of the experiments with a simulated beacon in Section 3.1 for the Optimal attack.

**Fig. D.2.**
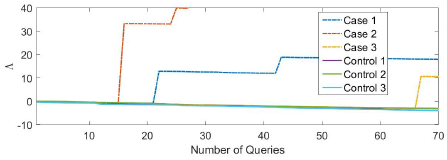
Example ;Ʌ distributions for 3 of the 40 case and 3 of the 20 control individuals of the experiments with a simulated beacon in Section 3.1 for the QI-attack.

### E Ground Truth for the Tests on Existing Beacons

In order to determine if the selected PGP(Personal Genomes Project) individual (PGP180/hu2D53F2) is in a beacon or not, we applied the SB attack on all the beacons used in Section 3.2. Thus, the decision made by the SB attack is independently used as the ground truth. The null hypothesis (the individual is not in the beacon) is rejected if *p* value is smaller than 0.05. The *p* value is calculated as *P* (*x ≥ k*; *x binomial*(*n,* 1 *- D*_*N*_))). Here, *N* is the size of the beacon, *k* is the number of “yes” responses to *n* asked queries, *x* is the response and *D*_*N*_ is the probability of no individual in the beacon having the queried allele. The tested individual had *p* value = 1 for all beacons and we concluded that s/he is not a member of any of the beacons we tested. In addition, the meta-data of the Kaviar beacon does not list this person as a member.

### F Experiments on Existing Beacons

Here, we show the evaluation results of our tests on the beacons of the beacon-network when an empty answer is not treated as a “no“. The results are shown in Table F.1. In summary, We could not detect the correct membership status for only 1 of the 9 beacons. The main difference of this case from the case in which an empty answer is treated as a “no” is that if empty answers are considered as a “no” response, the number of queries needed to determine beacon membership decreases significantly for some beacons. Nevertheless, if the empty answers are a result of two different genome copies (one in the beacon and one at hand) this conclusion would be incorrect.

**Table F.1.**
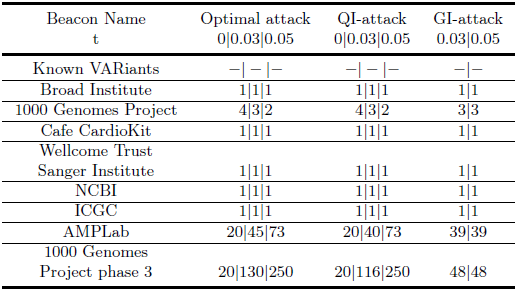
Number of queries required to receive a “no” within 1000 queries to existing beacons using an individual from PGP [2] when *t* = {0, 0.03, 0.05} for the Optimal attack [10], the QI-attack, and the GI-attack. Here, empty answers (i.e., the beacon has no information about the queried locus in the underlying dataset and returns neither a “no” nor a “yes“) are considered as a “no” response. “-” means in no “no” was found in 1000 queries.

### G GI-Attack without Population Information

In this section, we used a different training dataset to train the high-order Markov chain. The case and control individuals are the same as in the results shown in Section 3.1, that is from the CEU population. The high-order Markov chain was trained on the 77 individuals from the HapMap dataset “Mexican ancestry in Los Angeles” (MEX).

We observed that the GI-attack still performs very well, even when the population of the training dataset for the high-order Markov chain does not match the victim’s population as shown in Figure G.1.

**Fig. G.1.**
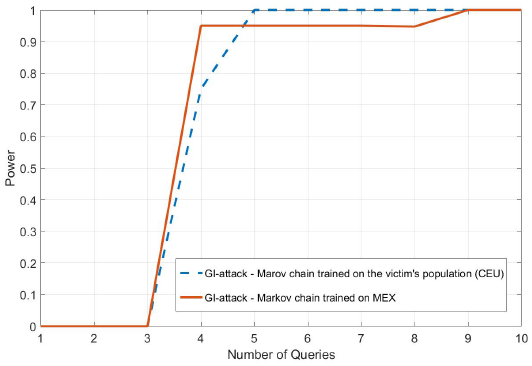
The GI-attack for *t* = 0.03 with the high-order Markov chain trained on the victim’s population (CEU) in comparison to the high-order Markov chain trained on a different population (here MEX) from the HapMap dataset.

### Footnotes

1 Linkage disequilibrium (LD) is a measurement for SNP correlations that shows which SNP positions are likely to be inherited together.

2 In this case “full” means that part of the DNA of the victim (e.g. a chromosome) is available in full and no locus is systematically hidden.

3 Such publicly available genome datasets are typically available with the population information about its anonymized participants. In such a case, we use a dataset that is consistent with the victim’s population to build our high order model. If the population information is not available in a dataset, it can be extracted by using ancestry inference techniques.

4 http://www.beacon-networg.org.

## References

1. Al Aziz, M.M., Ghasemi, R., Waliullah, M., Mohammed, N.: Aftermath of bustamante attack on genomic beacon service. BMC Medical Genomics 10(2), 43 (2017)

2. Church, G.M.: The personal genome project. Molecular systems biology 1(1) (2005)

3. Clayton, D.: On inferring presence of an individual in a mixture: a bayesian approach. Biostatistics p. kxq035 (2010)

4. Collins, F.S., Varmus, H.: A new initiative on precision medicine. New England Journal of Medicine 372(9), 793–795 (2015)

5. Gibbs, R.A., Belmont, J.W., Hardenbol, P., Willis, T.D., Yu, F., Yang, H., Ch’ang, L.Y., Huang, W., Liu, B., Shen, Y., et al.: The international hapmap project. Nature 426(6968), 789–796 (2003)

6. Homer, N., Szelinger, S., Redman, M., Duggan, D., Tembe, W., Muehling, J., Pearson, J.V., Stephan, D.A., Nelson, S.F., Craig, D.W.: Resolving individuals contributing trace amounts of dna to highly complex mixtures using high-density snp genotyping microarrays. PLoS Genet 4(8), e1000167 (2008)

7. Humbert, M., Ayday, E., Hubaux, J.P., Telenti, A.: Addressing the concerns of the lacks family: quantication of kin genomic privacy. In: Proceedings of the 2013 ACM SIGSAC conference on Computer & communications security. pp.1141–1152. ACM (2013)

8. Jacobs, K.B., Yeager, M., Wacholder, S., Craig, D., Kraft, P., Hunter, D.J., Paschal, J., Manolio, T.A., Tucker, M., Hoover, R.N., et al.: A new statistic and its power to infer membership in a genome-wide association study using genotype frequencies. Nature genetics 41(11), 1253–1257 (2009)

9. Ledford, H.: Astrazeneca launches project to sequence 2 million genomes. Nature 532(7600), 427 (2016)

10. Raisaro, J.L., Tramr, F., Zhanglong, J., Bu, D., Zhao, Y., Carey, K., Lloyd, D., Soa, H., Baker, D., Flicek, P., Shringarpure, S.S., Bustamante, C.D., Wang, S., Jiang, X., Ohno-Machado, L., Tang, H., Wang, X., Hubaux, J.P.: Addressing beacon re-identication attacks: Quantication and mitigation of privacy risks. The Journal of the American Medical Informatics Association 1(1), 1–1 (2016)

11. Samani, S.S., Huang, Z., Ayday, E., Elliot, M., Fellay, J., Hubaux, J.P., Kutalik, Z.: Quantifying genomic privacy via inference attack with high-order snv correlations. In: Security and Privacy Workshops (SPW), 2015 IEEE. pp. 32–40. IEEE (2015)

12. Sankararaman, S., Obozinski, G., Jordan, M.I., Halperin, E.: Genomic privacy and limits of individual detection in a pool. Nature genetics 41(9), 965–967 (2009)

13. Shringarpure, S.S., Bustamante, C.D.: Privacy risks from genomic data-sharing beacons. The American Journal of Human Genetics 97(5), 631–646 (2015)

14. Siva, N.: 1000 genomes project. Nature biotechnology 26(3), 256–256 (2008)

15. Visscher, P.M., Hill, W.G.: The limits of individual identication from sample allele frequencies: theory and statistical analysis. PLoS Genet 5(10), e1000628 (2009)

16. Wang, R., Li, Y.F., Wang, X., Tang, H., Zhou, X.: Learning your identity and disease from research papers: information leaks in genome wide association study. In: Proceedings of the 16th ACM conference on Computer and communications security. pp. 534–544. ACM (2009)

